# Genomic prediction of single cross families of perennial ryegrass in two nitrogen managements

**DOI:** 10.64898/2026.05.05.722839

**Authors:** Divino Rosa dos Santos Junior, Dario Fè, Ingo Lenk, Christian S. Jensen, Torben Asp, Luc Janss, Elesandro Bornhofen

**Affiliations:** Center for Quantitative Genetics and Genomics, Aarhus Univ., Aarhus, Denmark; Research Division, DLF Seeds A/S, Store Heddinge, Denmark

**Keywords:** Genomic selection, genotype-by-environment interaction, non-additive effects, GWAS

## Abstract

The performance of a single cross is determined by the average additive effects of the parents, as well as the interactions between them. These quantities can be estimated using an appropriate genetic design, allowing for the estimation of general (GCA) and specific (SCA) combining abilities. The prediction of GCA for new parents and the total genetic value of unrealized crosses can be made when genome-wide marker information is available. Several studies in crops such as maize and rice have demonstrated the potential of genomic-assisted prediction of single-cross performance in economically important crops. However, no study to date has explored its relevance in perennial ryegrass, an obligate allogamous species that is bred in genetically heterogeneous families. In this study, we aimed to estimate genetic parameters and assess the ability of genomic models to predict the performance of F2 families in terms of dry matter yield and nutritive quality traits. We used data from a large partial diallel involving 104 parents from two distinct subpopulations, as inferred by admixture analysis. F2 families were evaluated in multiple environments and under two nitrogen availability conditions. Genotyping-by-sequencing of the parent plants produced 42,145 variants after quality control, which were used to estimate genomic relationships based on identity-by-state. Variance component estimation revealed limited GCA and SCA interactions with the environment, and particularly with nitrogen management. The predictive abilities of two parental models exceeded 0.60 and often surpassed 0.70 for most traits. However, incorporating non-additive effects into the model did not improve predictive ability. We leveraged the genetic diversity among parents to map genomic regions associated with all recorded traits. Genome-wide association studies (GWAS) by genomic best linear unbiased prediction (GBLUP) identified six quantitative trait loci (QTL) regions, with 45 candidate genes within the linkage disequilibrium range, estimated at approximately 92 kb. Our results demonstrate that genomic prediction of single crosses can be performed with high accuracy, especially when both parents are also progenitors of families in the training set.

## Introduction

Perennial ryegrass (*Lolium perenne* L.) is a forage crop of global importance, widely distributed in temperate regions of the Americas, Asia, and Europe (Casler et al., 2025). Its large-scale production is justified by its frequent application in animal feed, due to its excellent nutritional quality and high potential for dry matter yield (DMY). However, both nutritional quality and DMY are intrinsically linked to the adequate availability of nitrogen (N), which plays a key role in increasing crude protein content and influencing fiber composition and digestibility (Cardenas et al., 2019; Sandaña et al., 2021). In this context, considerable attention is directed towards the adverse effects of applying nitrogen fertilizers in agricultural crops (Firoozzare & Naghavi, 2021), especially in countries like Denmark, where farmers are subject to strict guidelines and laws regulating the use of N (Blicher-Mathiesen et al., 2015; Hoffmann et al., 2020). This highlights the importance of modern pasture production systems facing the challenge of improving N fertilization management. The goal is to reduce N rates while increasing forage yield, resulting in greater N use efficiency (NUE, forage yield per unit of applied N) (Liu et al., 2021). A promising approach is the development of cultivars with enhanced nitrogen uptake and utilization efficiency, capable of maintaining vigorous vegetative growth, improving crop yields, and converting nitrogen into protein more effectively, which would lead to lower production costs and reduced environmental impact (Fischer & Connor, 2018; Peters et al., 2021; Langworthy et al., 2023). Such cultivars are particularly valuable in nitrogen-limited conditions, where efficient nitrogen use is essential for maintaining high performance in suboptimal environments. Achieving such improvements, however, requires breeding strategies capable of accelerating genetic gain.

A shift of breeding goal towards NUE would require a considerable amount of time before any noticeable impact at farm level. Genomic selection has been used to shorten breeding cycles across different crop species. The method uses statistical models coupled with genome-wide markers to forecast the genetic value or model genomic relationships in a training population with phenotypic and genotypic data (Meuwissen et al., 2001). Since its integration into plant breeding, numerous methodologies and statistical models have emerged to apply genomic prediction (GP) in both annual and perennial plant species, aimed at expediting genetic progress. In perennial ryegrass, genomic selection models have been developed and validated for many complex traits such as DMY (Pembleton et al., 2018; Arojju et al., 2020), nutritive quality traits (Arojju et al., 2020; Malmberg et al., 2023) and heading date (Fè et al., 2015). Genomic selection in perennial ryegrass has also been explored under varying nitrogen conditions, with Zhao et al. (2020) using a collection of 184 global accessions of ryegrass and demonstrating comparable accuracy in predicting traits under normal or low-N treatments, with notably moderate prediction accuracies observed for NUE under low-N stress, emphasizing the necessity for future research to enhance the prediction accuracy of NUE through larger sample sizes and increased SNP markers in field-grown plants under N-deficient conditions.

In addition to predicting the genetic merit of untested individuals, GP models can be used to guide crosses in breeding programs by predicting the general combining ability (GCA) of available parents and the specific combining ability (SCA) of potential parental combinations. GCA measures the reproductive value of parents across multiple crosses and is essential for predicting the medium- and long-term response to selection. On the other hand, SCA captures non-additive effects, which significantly influence the performance of offspring in specific crosses (Sprague & Tatum, 1942). GP models for hybrids, particularly in crops like maize and sorghum, have shown that including GCA and SCA improves predictive accuracy, enabling more precise estimates of genetic effects (Fonseca et al., 2021; Sapkota et al., 2023). However, despite the well-established importance of these parameters in maximizing selection gains, there is still limited information on their inclusion in segregating populations of forage species such as ryegrass (Zhao et al., 2020).

Besides using all genome-wide variants simultaneously for prediction tasks, single variant regression known as genome-wide association studies (GWAS) can reveal genomic regions linked to the expression of quantitative traits (QTLs). It has been applied in ryegrass to identify variants related to traits such as dry matter yield, nutritive quality, and stress resistance (Pembleton et al., 2018; Jaškūnė et al., 2020; Colas et al., 2022). Overall, this approach holds great potential for dissecting the genetic architecture of complex traits, providing insights into genomic regions associated with traits of interest. In forage breeding programs aiming to improve complex traits such as biomass production and nutritive quality, this information can support more efficient breeding strategies by enabling the identification of useful QTLs and marker-assisted selection. Thus, GWAS provides complementary information to genomic prediction by enabling the identification of genomic regions underlying trait variation, which can support marker-assisted selection and biological interpretation.

The present study leverages a large sparse diallel of perennial ryegrass tested in two nitrogen conditions across multiple environments to estimate genetic parameters and compare parental models in terms of predictive ability. In this context, and to our knowledge, this is the first study to apply genomic prediction of single-cross performance in perennial ryegrass using a GCA/SCA framework. Models are also compared under varying levels of relatedness between training and testing sets. The goal is to demonstrate the utility of modelling GCA and SCA for full-sib family prediction under varying N availability and to investigate genomic regions associated with DMY and nutritive quality traits.

## Material and methods

### Plant material and Experimental Design

This study used data of full-sibs families and respective parental single-plants of perennial ryegrass, previously published by (Bornhofen et al., 2022) for genome-wide analysis. The families were developed using a partial diallel mating design comprising 104 parents, chosen to represent a wide range of the genetic spectrum used in commercial breeding. A total of 381 families were obtained through two disconnected partial diallels: the larger diallel (Diallel A) included 88 parents, resulting in 335 crosses, while the smaller one (Diallel B) involved 16 parents, yielding 46 crosses. Parental plants were clonally propagated to ensure consistent genetic material across multiple crosses. First generation seeds (consisting of full-sibs) were sown in isolated field plots for seed multiplication. Second generation seeds were sown in the experiment.

The experiment was conducted at two locations over two testing years, with four harvests of fresh biomass performed per year. Families were randomly assigned to two nitrogen conditions (N) – Normal (380 kg N ha^−1^ yr^−1^) and Low (280 kg N ha^−1^ yr^−1^). Therefore, a total of 1,024 experimental units, each measuring 12.5 m^2^ (1.5 m wide by 8.35 m in length) were evaluated in location 1 (Store Heddinge, DK). These plots were arranged in an irregular rectangular grid of 144 rows by eight columns. The same setup was defined for location 2 (Skælskør, DK) however, the plot size was larger (13.5 m2; 1.5 by 9 m), and the trial was organized in a regular rectangular grid of 128 rows by eight columns. Each field experiment layout consisted of entries randomly assigned to one (254 families plus four controls) or two (127) replicates within nitrogen availability conditions: normal N rate and low N rate.

Regarding nitrogen management, two different application strategies were implemented based on the availability of nitrogen. Under normal conditions, nitrogen was applied at rates of 152 kg N ha^−^ at the onset of spring growth, followed by 114 kg N ha^−^ after the first cut, 76 kg N ha^−^ after the second cut, and 38 kg N ha^−^ after the third cut. For the low nitrogen availability treatment, the application rates were adjusted to 112 kg N ha^−^ at the start of spring growth, 84 kg N ha^−^ after the first cut, 56 kg N ha^−^ after the second cut, and 28 kg N ha^−^ after the third cut. Additionally, irrigation was provided immediately following each nitrogen fertilization event.

### Determination biomass yield and nutritive quality traits

The harvests were carried out at approximately 5-week intervals, depending on the meteorological conditions. The first harvest took place in the spring of 2019, when the plants reached the boot (R0) growth stage (Moore et al., 1991). The harvests were conducted using a HALDRUP plot combine, with four fresh biomass cuts mechanically executed each year, covering two testing years (2019 and 2020). The dry matter yield refers to the total dry biomass harvested over the year, i.e., the sum of the four cuts made during the vegetative period. Nutritive quality was estimated using a near-infrared reflectance spectroscopy (NIR) device installed on the combine, which scanned the sample at each harvest. Quality traits were predicted using calibration models and calculated as weighted averages, with each harvest weighted according to its contribution to the total annual biomass. The traits included acid detergent fiber (ADF), acid detergent lignin (ADL), dry matter digestibility (DMDig), neutral detergent fiber (NDF), digestible NDF (NDFD), protein (Prot), and water-soluble carbohydrates (WSC). All quality parameters were expressed as a percentage of DMY, except for NDFD, which was expressed as a proportion of NDF.

### Genomic data

The genotyping of single plants used as parents in the diallel was performed via genotyping-by-sequence, yielding 42,145 variants (Bornhofen et al., 2022). Variants were filtered using BCFtools v1.17 (Danecek et al., 2021) to retain sites with a minimum quality score of 20 (QUAL ≥ 20), a minimum mapping quality of 10 (MQ ≥ 10), minor allele frequency ≥ 0.05, and missingness ≤ 50%, after which genotype imputation was performed using Beagle v5.5 (Browning et al., 2018).

## Statistical analysis

### Model fitting

Response variables were modeled as a function of different sources of variation in a linear mixed model (LMM) framework using the software DMU (Madsen & Jensen, 2013) . We overlaid design matrices of parents 1 and 2 for the estimation of main effect and interactions as they appear in crosses both as female and male plants. For each trait, the following two LMM models were fitted to the data:

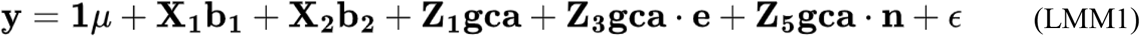

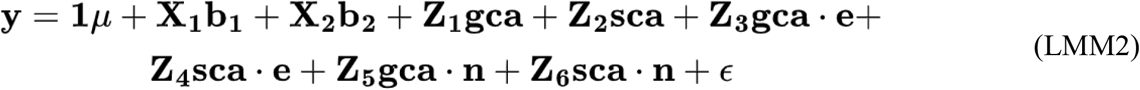

where, is the vector of phenotypes; represents the overall intercept with associated vector containing 1’s; is a vector of fixed effects of environment and nitrogen management classes, is the fixed effect for classes created by combine harvesters used and the date of harvest as we noticed a substantial systematic field variation generated by such classes; and are the associated incidence matrices for the fixed effects, respectively; is a vector of random parental GCA effects assumed 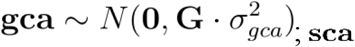 is vector of random SCA effects for parental combinations assumed 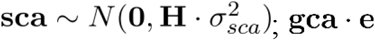 is a vector of random GCA-by-environment interaction effects, following 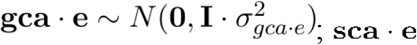 is a vector of the random interaction effect of SCA-by-environment, assumed 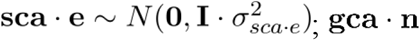 is a vector for the random interaction effects between GCA and the nitrogen management,following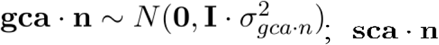 is a vector of random SCA by nitrogen management effects, assumed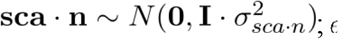 is the vector of random residual effect, which follows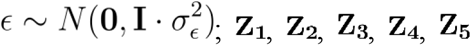, **Z**_6_ are incidence matrices linking phenotypic observations to the respective random effects, and refers to an identity matrix. The additive genomic relationship matrix **G** for parents was computed according to method 1 (VanRaden, 2008), following the expression: **G =WW′/**2Σ*p*_*i*_(1− *p*_*i*_), where is a matrix of centered genotypes at each marker. The covariance matrix for SCA was obtained by the Kronecker product of with itself:**H**=**G**⨂**G**. The genetic value of each family was computed as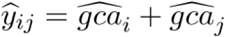 for LMM1 and 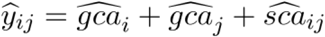for LMM2. The (co)variance components of LMM2, estimated using restricted maximum likelihood (REML), were used to calculate genetic parameters of interest. The phenotypic variance was obtained by the expression:

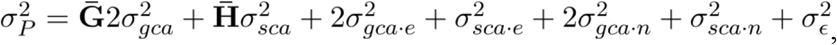

where 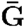and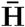 are the average of the diagonal elements of matrices **G** and **H**, respectively. The broad-sense heritability at the plot level was computed as 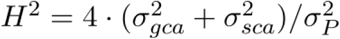and the plot-level narrow-sense heritability as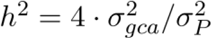.

### Evaluation of model performance

The ability of both LMM models to predict family performance was assessed via 5-fold cross-validation (CV) repeated 10 times. Cross-validation was executed using the DMU4 module of the DMU software (Madsen & Jensen, 2013) using variance components estimated in the parental model described before as priors. For each run of the 5-fold CV, the 381 families were randomly allocated to five groups. Therefore, the training set was composed of 80% of the families, while 20% were allocated to the testing set. Prediction accuracy was measured as the Pearson correlation coefficient between predicted values and mean genotypic values adjusted for fixed effects present in the parental model. We also investigated the impact of relatedness on prediction accuracy by three different random cross-validation scenarios (Figure 1). In T2, we sampled families to compose the testing set with the constraint that all parents of families in the testing set were in common with those from the training set. For the remaining CV schemes, we removed from the training set all family records of either one (T1) or both (T0) parents of families in the testing set. In all three CV scenarios, the training set size was adjusted to 200 families, a constraint driven by T0. The process of sampling families for allocation to the training set and the masking of records of parents in the training set was repeated 25 times.

**Figure 1.**
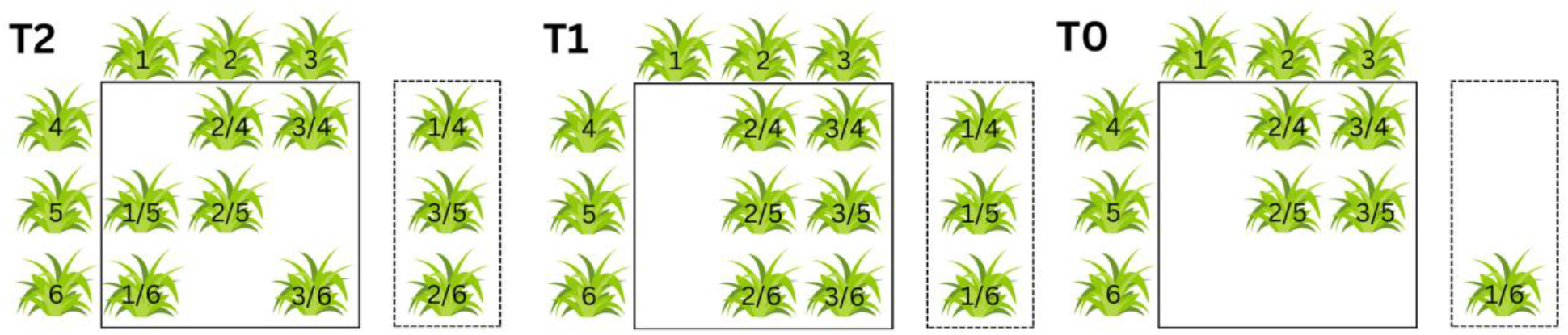
Schematic representation of the three cross-validation scenarios used to evaluate the effect of the relationship on prediction accuracy. Ryegrass families within solid lines compose the training set while those within the dashed lines represent the testing set. In T2, both parents of each family in the testing set were also used in crosses of families of the training set. In T1, only one of the parents of families in the testing set is also one of the parents of families in the training set. Finally, no parental genotype is shared between training and testing sets in T0.

### Genome-wide association analysis

We carried out GWAS by GBLUP where SNP effects 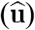 were obtained by back-solving the solutions for the GCA estimated with model LMM2, according to the following equation:

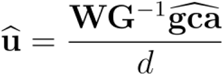

where,**W** is the allele frequency corrected marker matrix obtained as **W =M**−2* (*p*−0.5), where contains SNP markers coded as -1, 0, and 1; is the inverse of the genomic relationship matrix (**G**) described before;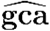 is the vector of general combining ability values of parents; and is a scaling factor calculated as 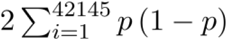, where *p* is the allele frequency of the marker, and 42145 is the number of variants. The approximated p-values for marker effects were obtained according to a two-tailed hypothesis test based on a t-distribution (Gondro, 2015; Waters et al., 2022). A Bonferroni correction of 5% (0.05/42145) was used to declare significant SNPs, which were clumped to remove significant variants in LD with the leading SNP at each peak region. Finally, candidate genes were identified by examining the region surrounding significant markers according to the estimated LD decay. Functional annotation of candidate genes was performed using PANNZER2 (Törönen et al., 2018) with protein sequences retrieved from the reference genome (Nagy et al., 2022).

## Results

The effect of nitrogen availability management was small for most of the traits across environments, except for dry matter yield (Figure 2A), which showed an increase of 10– 20% under normal nitrogen conditions compared to low nitrogen depending on the harvest. In contrast, the remaining traits exhibited only minor differences between nitrogen levels. Empirical BLUEs from model LMM1 show a greater influence of year–location combinations on family performance across all measured traits These traits are not independent, and often high correlations are reported among them. We used genotypic values adjusted for fixed effects present in the parental models to compute simple and partial correlations, which is relevant for comparison of variance components and predictive abilities across traits explored later. As expected, fiber related traits, digestibility, and water soluble carbohydrates were highly correlated among them (Figure 2B). Simple pairwise correlations also revealed small to moderate correlations between dry matter yield and nutritive quality traits. However, most of these correlations disappeared when partial correlations were computed, revealing two moderate values between ADL and DMDig (−0.65) and NDF and ADF (0.66). A small partial correlation of -0.37 was observed between protein content and water soluble carbohydrates.

**Figure 2.**
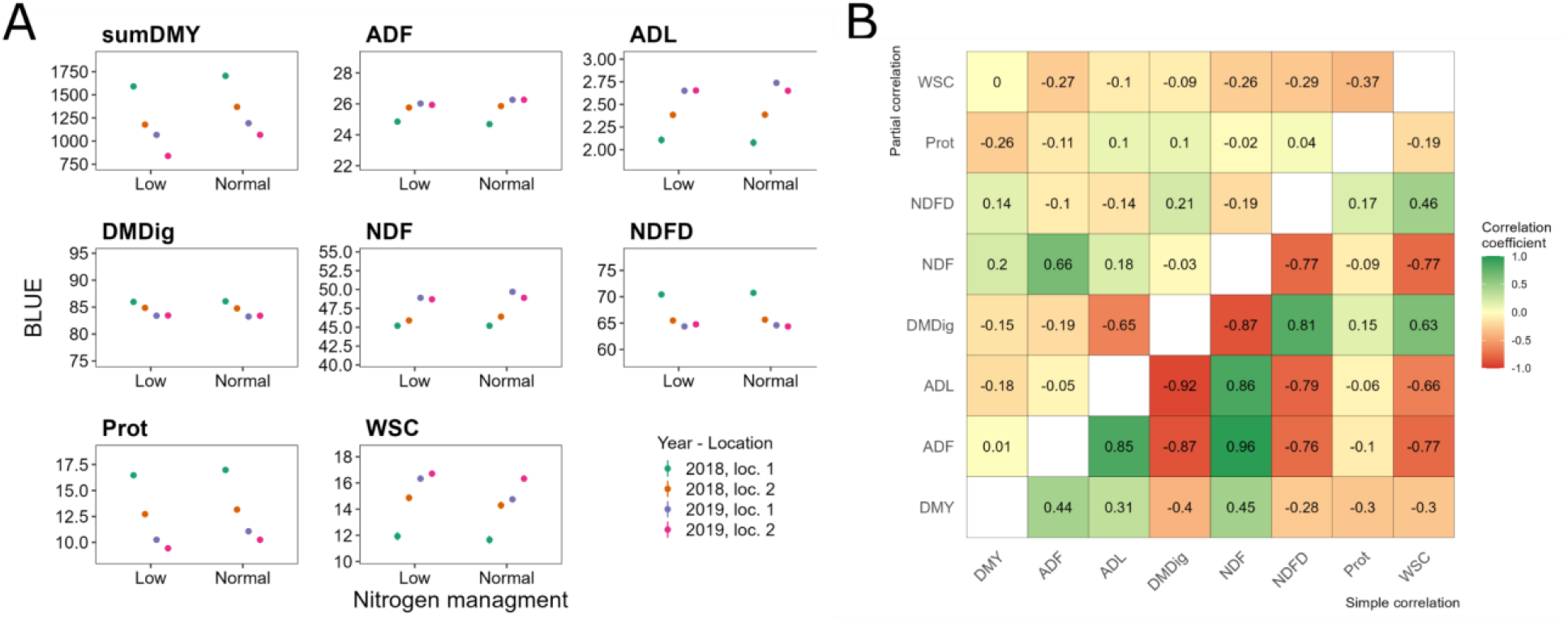
Environmental effects on traits were mostly driven by year and location rather than nitrogen management. **A:** Empirical Best Linear Unbiased Estimators (BLUEs) of environments for eight traits in two nitrogen availability scenarios (low and normal). **B:** Simple (lower triangle) and partial (upper triangle) correlation coefficients between all pairs of traits recorded on 381 families of perennial ryegrass assessed in multiple locations. Correlations were computed with adjusted means.

### Population structure of parental genotypes

The individual admixture proportions for a model with the value of K = 2, which was estimated through cross-validation, revealed the ancestral composition of each parent in the diallel (Figure 3A). The vertical bars represent this ancestral composition and are sorted according to subpopulation membership. The existence of two subpopulations, and some extent of admixture between them, is also suggested by inspecting the scatter plot of the principal component analysis (Figure 3B). Additionally, members of subpopulation SP1 are shown more scattered, suggesting higher genetic diversity within that group. The extent of genetic diversity for the whole set of parents can be inferred by the rapid LD decay of 92.11 Kb distance to reach half of its maximum (Figure 3C), due to high recombination rates in the population from where the parents belong, which is expected due to the allogamous nature of the species. Finally, a table with all the realized crosses is depicted in Figure 3D. Colors indicated a well balanced proportion of crosses made between parents both within and between sub-populations.

**Figure 3.**
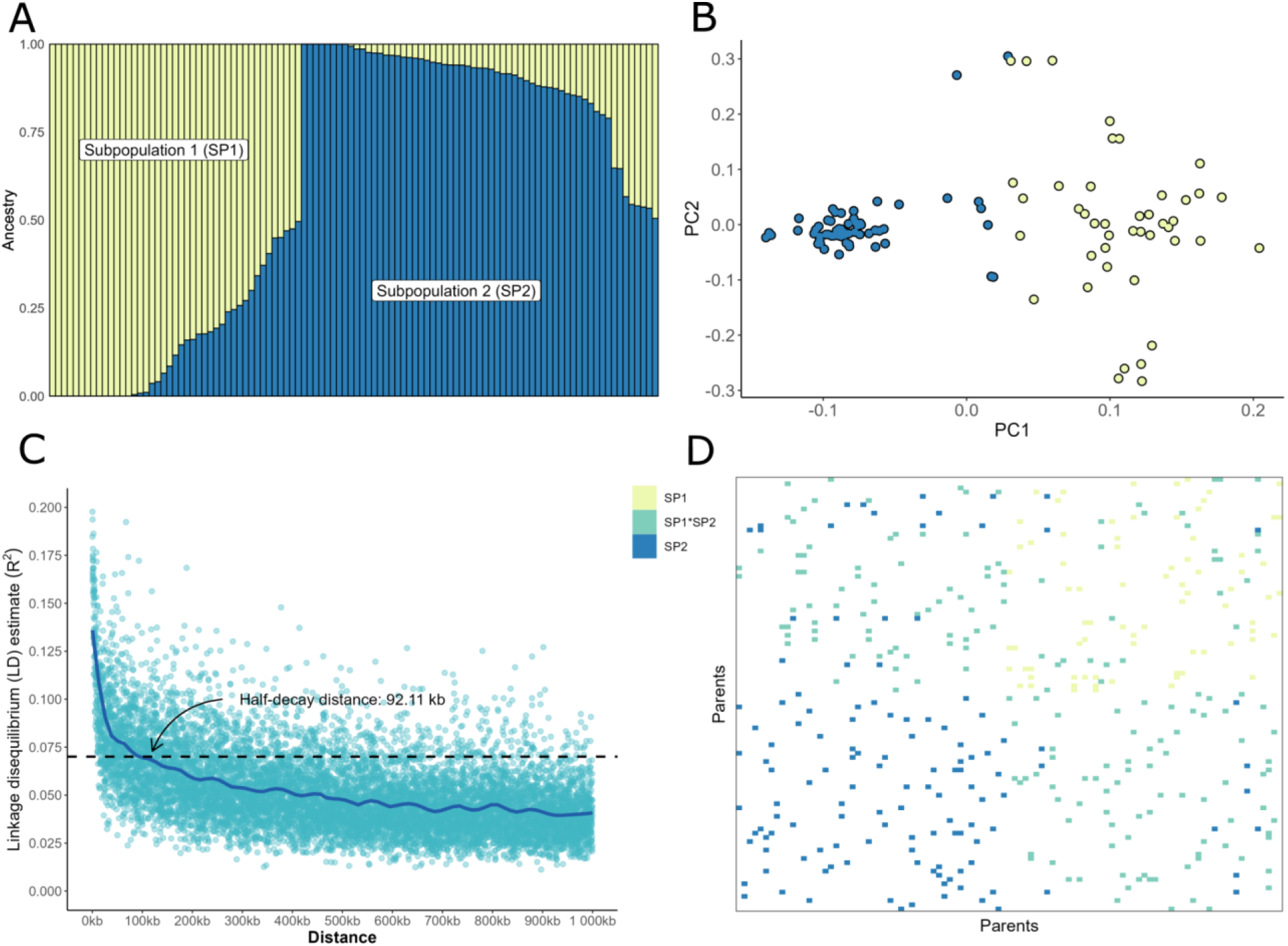
Diallel parents belong to two perennial ryegrass subpopulations. **A:** individual admixture proportions for a model with K equals to 2, estimated via cross-validation. Vertical bars represent the ancestral composition of each parent, and are sorted according to subpopulation membership. **B:** scatter plot depicting the first two principal components computed from SNP markers. Each point in the graph represents one of the 104 parents and is colored according to the ADMIXTURE results in A. **C:** linkage disequilibrium (LD) according to physical distance. **D:** partial diallel structure with tiles representing each of the 381 crosses and colored according to the subpopulation origin of the parents (SP1: both parents belong to subpopulation 1, SP2: both parents belong to subpopulation 2, and SP1*SP2: crosses between parents from different subpopulations). Parents on x- and y-axis are sorted according to their genetic similarity obtained by hierarchical clustering analysis with the complete agglomeration method. The dissimilarity metric used was 1 minus the genomic correlation, obtained by calling the “cov2cor” R function on the genomic relationship (**G**) matrix.

### variance components and heritabilities

Heritability estimates were moderately high (0.6–0.79) for most evaluated traits (Figure 4), except for NDFD, Prot, and WSC, which showed low heritability values (0–0.4), indicating a stronger environmental influence on their phenotypic expression. Across all evaluated traits, broad-sense heritability was higher than narrow-sense heritability, indicating that additive genetic variance represents a major component of the total genetic variation. However, for DMY the estimates of broad- and narrow-sense heritability were more similar, indicating that non-additive genetic effects may also contribute to the expression of this trait.

**Figure 4.**
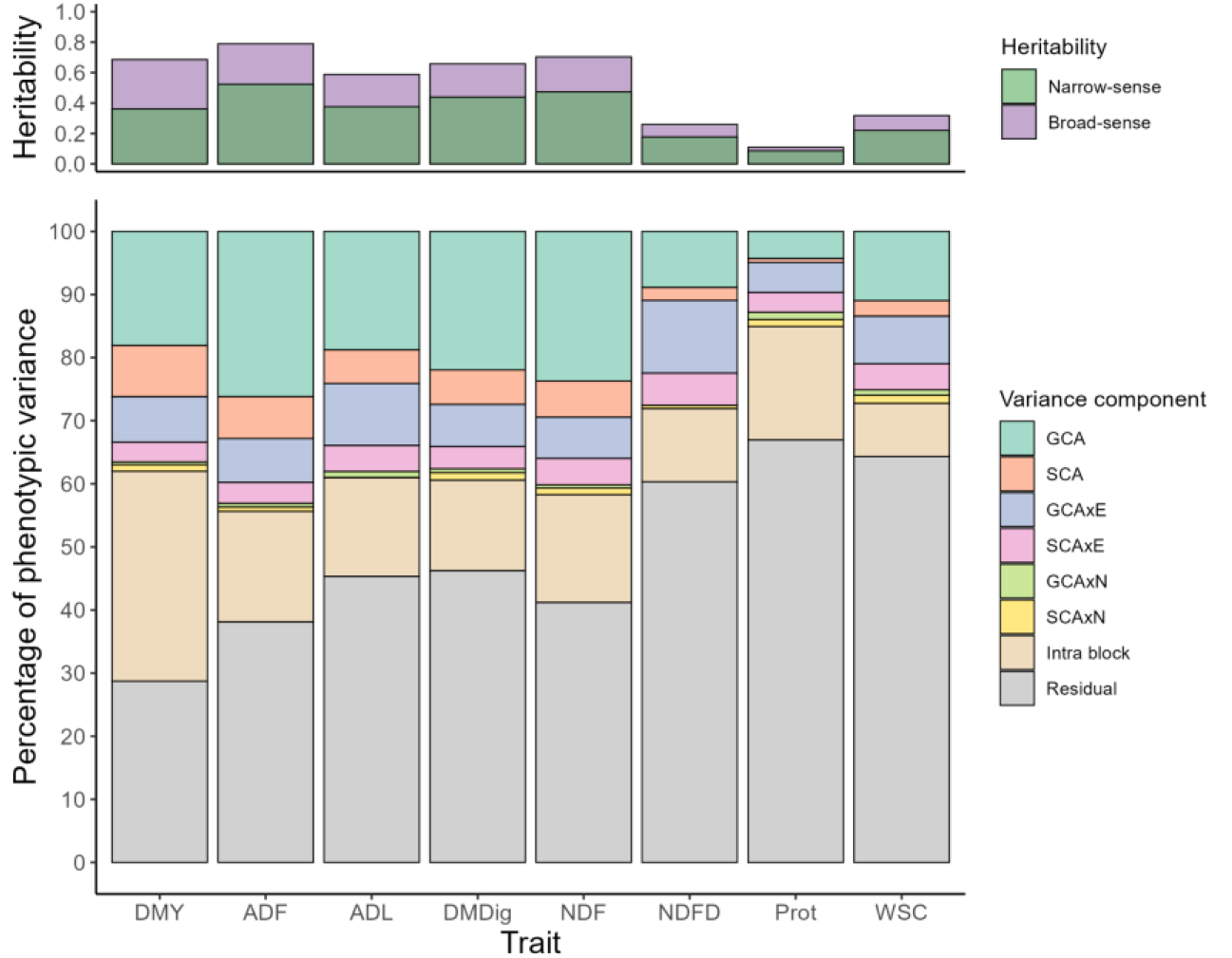
Causal variance decomposition reveals the importance of additive genetic effects in the performance of F2 families of ryegrass. The upper bar plot depicts the proportion of the phenotypic variance that is due to additive genetic effects (narrow-sense heritability) and total genetic effect (broad-sense heritability) at the plot level. The bottom bar plot reveals the distribution of the observational components of variance, where GCA is the general combining ability, SCA is the specific combining ability, E refers to environments, and N to the nitrogen management effect.

The analysis of the general combining ability (GCA) of each parent and the specific combining ability (SCA) between parent pairs revealed clear differences in genetic effects. GCA captures additive genetic effects across multiple crosses, whereas SCA reflects the specific performance of a cross, accounting for non-additive interactions and parental allele frequency differences (Hallauer et al., 2010). Partitioning of trait variation revealed a predominance of additive genetic effects, as expected, while a non-negligible proportion of the variance was attributable to non-additive genetic effects captured by the SCA term. This observation prompted the question of whether incorporating non-additive effects would translate into higher prediction accuracies in genomic selection for perennial ryegrass, which is addressed in the subsequent section.

The interactions between GCA and SCA with the environment were generally lower than GCA and SCA, with the exception of NDFD and protein content, suggesting that genetic effects were relatively stable across environments. In contrast, interaction effects with nitrogen management were negligible, with both GCA × N and SCA × N components contributing close to zero to the total phenotypic variance. This indicates that nitrogen management had little influence on the contribution of combining ability effects, as the interaction components explained only a small proportion of the phenotypic variance. Consequently, the relative contribution of additive (GCA) and non-additive (SCA) genetic effects remained largely unchanged across nitrogen regimes for forage quality traits and dry matter yield in ryegrass. Residual and spatial variation, captured by the intra-block component of the model, accounted for the majority of the phenotypic variance.

### Prediction accuracy of single-cross families

We compared two linear mixed models for their ability to predict untested crosses. The two models, named LMM1 and LMM2, differed by the inclusion or not of the SCA component. Therefore, we aimed to assess the extent to which non-additive effects play an important role in the performance of families. For most of the traits analyzed, the predictive accuracy was above 0.7, indicating, in general, a high level of predictive accuracy (Figure 5). This result may be partly explained by the high precision of the adjusted family means used as validation targets. The large number of observations per family reduces sampling error in phenotypic estimates, which can inflate the observed prediction accuracy. Furthermore, the inclusion of non-additive effects in the model did not result in significant gains in accuracy despite a relevant SCA effect for some of the traits (Figure 4).

**Figure 5.**
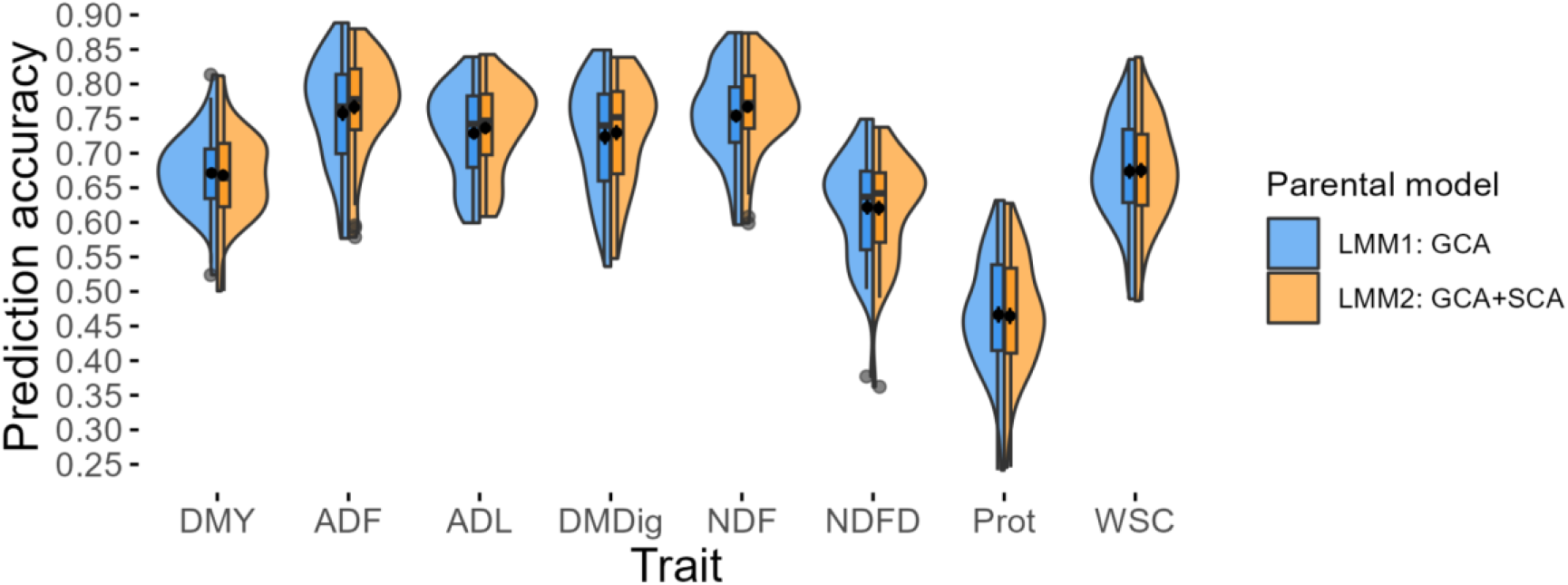
Prediction accuracies remained mostly unchanged despite the inclusion of the specific combining ability effect. Accuracies were computed as the Pearson correlation coefficient between predicted family genetic values from a repeated 5-fold CV scheme and adjusted means. GCA and SCA are the general and specific combining ability, respectively.

Since genotyping data for the families were available (Bornhofen et al., 2022), we compared the prediction accuracy of LMM1 with that of a GBLUP model (named progeny model) that uses a genomic relationship matrix **G** computed from allele frequencies of genome-wide variants within the families themselves (see Ashraf et al., 2014; Cericola et al., 2018 for details on the **G** computation). The results, shown in Supplementary Note S1, reveal an average prediction accuracy that is 10% higher across traits for the progeny model; however, this improvement comes at the expense of requiring nearly four times more genotyped samples. In addition, parental genotypes are available earlier than those of the F2 progeny, which has important implications for early selection and breeding decision-making.

### Prediction accuracy under varying levels of relatedness

Prediction accuracy can be significantly influenced by family structure and the degree of relatedness between individuals. To assess this effect, we tested three different validation schemes. Additionally, we evaluated the impact of incorporating non-additive effects, such as SCA, on prediction accuracy. Our results indicate that, regardless of the validation scheme used, including SCA did not lead to a significant improvement in predictive accuracy. In contrast, increasing the level of relatedness among individuals in the training and testing sets increased predictive accuracy. Predictive accuracy for dry matter yield increased by approximately 70–80% when comparing the most restrictive scenario (T0) with the scenario where both parents were present in the training set (T2). However, even under the most restrictive validation scheme (T0), where the parents of the crosses in the test set were not included in any of the crosses in the training set, we still observed predictive accuracies above 0.4–0.5 for most traits despite the broad diversity of the selected parents (Figure 6).

**Figure 6.**
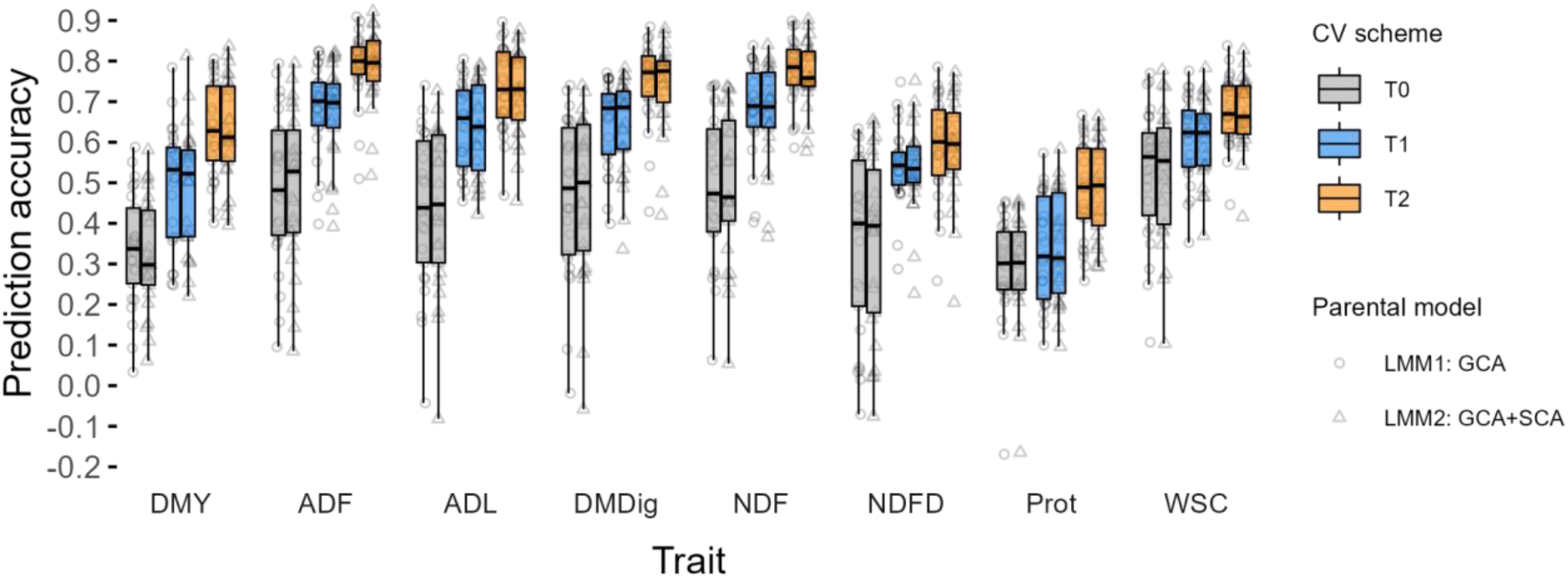
The training set composition has a large effect on the ability to predict F2 family performance in perennial ryegrass. Prediction accuracy was estimated by random cross-validation with varying degrees of relatedness between training and testing sets. In T0, none of the parents are common between training and testing sets while for T1 and T2, one or both parents, respectively, are shared.

### Genome-wide association analysis

We conducted genome-wide association studies (GWAS) using GBLUP to leverage the high diversity of the diallel parents, which is an essential requirement for GWAS. Marker-trait associations (MTAs) were detected for all traits except DMY, totaling 23 MTAs across six genomic regions (Figure 7A), with 5 MTAs for NDFD and WSC, 4 for NDF, 3 for ADF, ADL, and DMDig each, and 2 for Prot. The clustering of MTAs reflects the high correlations among certain traits (Figure 2B), as genetically correlated traits may share underlying genetic architecture, leading to co-localization in the same genomic regions. For downstream analyses, we focused on the leading SNPs in each region. The prominent peak on chromosome 1 is led by SNP 1_249271039_G_A, located within the coding region of V3.Lp_chr1_0G18262, associated with gene regulation and developmental processes (Supplementary Table S1). Additive effects of the SNPs were illustrated with effect plots showing GCA values as a function of allele dosage (Figure 7C). SNP 3_312031270_T_C is linked to a gene encoding a protein with a 5’-3’ exonuclease domain, potentially influencing DNA repair or RNA degradation, while SNP 6_73570603_T_C is associated with a gene encoding the D-like subunit of Glutamyl-tRNA(Gln) amidotransferase, involved in glutamine and amino acid metabolism. Although principal components were not explicitly included, population structure was effectively controlled through the genomic relationship matrix in the GBLUP framework, thereby minimizing the risk of spurious associations due to stratification. Consequently, the detected MTAs are unlikely to reflect population structure artifacts. Figure 7B shows the genomic regions with nearby genes, additional SNPs, and linkage disequilibrium patterns as a heatmap.

**Figure 7.**
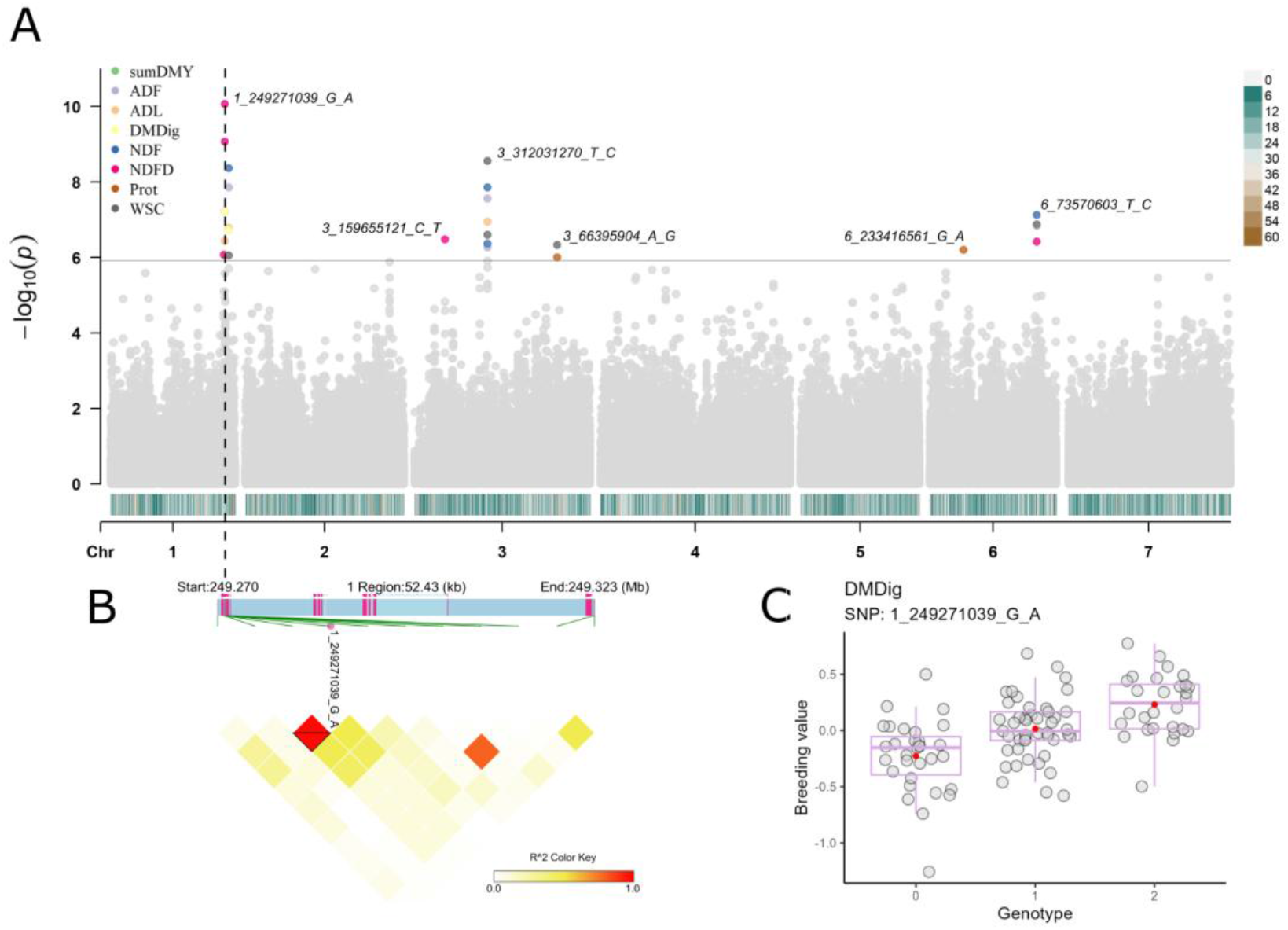
Genome-wide association analysis by GBLUP reveals six QTLs for nutritive quality traits in perennial ryegrass. **A:** Multi-trait Manhattan plot showing significant variants defined by the Bonferroni Threshold at 5%. The segment on the bottom of each chromosome represents SNP density for 2249 bins of 1 mega base (Mb) each. **B:** linkage disequilibrium (LD) heatmap displaying a region of 53.43 Kb where the leading SNP (1_249271039_G_A) is located. **C:** distribution of breeding values for DMDig according to the allele state.

## Discussion

In this study, genomic prediction proved to be a promising tool for predicting family performance in perennial ryegrass. Predictive abilities were moderate to high across traits, indicating that genomic models can effectively capture genetic variation under different nitrogen conditions. Breeding values from genome-wide statistical models can accelerate genetic gains, especially for traits that are difficult to measure, such as nitrogen use efficiency. Previous studies in perennial ryegrass have also reported successful prediction of traits such as heading date, resistance to crown rust, seed production, and nutritive quality (Fè et al., 2015, 2016; Guo et al., 2018; Pembleton et al., 2018; Konkolewska et al., 2023). Modelling the genetic effect of parents in addition to nitrogen availability has been explored in several crops, such as maize (Fristche-Neto et al., 2018; Costa-Neto et al., 2021; Fonseca et al., 2021), rice (Labroo et al., 2021), sorghum (Sapkota et al., 2023), but remains relatively unexplored in forage grasses, highlighting the relevance of the present study.

In the context of this study, although nitrogen availability affected some traits, particularly dry matter yield (DMY), no significant interaction between nitrogen management and combining abilities was detected. This indicates that the relative ranking of genotypes remained largely stable across nitrogen levels, with families performing well under high N also tending to perform well under low N. Similar patterns have been reported by Sapkota et al. (2023) in sorghum for several agronomic traits related to hybrid performance, whereas (Luz et al. (2024) observed significant GCA × N and SCA × N interactions for traits such as grain yield and plant architecture in maize under contrasting nitrogen regimes. In perennial ryegrass, nitrogen availability is known to influence several forage quality traits, including protein concentration, digestibility, and carbohydrate accumulation (Sandaña et al., 2021), while substantial genetic variation has been reported for these traits (Colas et al., 2022). The limited interaction observed may be influenced by the relatively narrow contrast between nitrogen treatments evaluated in this study. Future studies exploring a wider range of nitrogen environments may reveal interactions not detected here.

As genotype-by-environment and by nitrogen effects were small, we focused our genomic prediction study on predicting the average breeding or genetic values. Incorporating GCA and SCA components has been shown to increase prediction accuracy, especially in inter-heterotic crosses, where the greater genetic distance between parental groups enhances hybrid vigor (Ramstein et al., 2020; Costa-Neto et al., 2021; Rogers et al., 2021). In these cases, SCA plays a more significant role in improving hybrid prediction accuracy. In contrast, in crosses within the same heterotic group or with a smaller genetic distance, SCA tends to be less pronounced, and selection tends to be more influenced by the GCA effect (Albrecht et al., 2014; Fristche-Neto et al., 2018). Our results suggest no benefit in terms of prediction accuracy when non-additive genetic effects are included, at least considering the parametrization adopted here. While SCA effects did not improve prediction accuracy in this study, their contribution may depend on the experimental design and data structure, and thus should be interpreted with caution. Similar results were found by Labroo et al. (2021) in rice for hybrids aimed at increasing predictive accuracy, comparing models with only GCA to models that included both GCA and SCA, without observing significant differences between them. Similarly, Melchinger et al. (2023) simulated hybrid populations based on molecular data from two maize experimental datasets and developed GBLUP estimates for hybrids based on expected prediction accuracy and GCA and SCA effects, concluding that the prediction accuracy of hybrids largely depends on the GCA of their parental lines. These findings suggest that non-additive genetic effects may contribute less to predictive ability in these contexts, or may be partially captured as additive effects(Huang & Mackay, 2016; Vitezica et al., 2017), aligning with the results of this study.

The degree of relatedness between the training and validation sets is a major determinant of prediction accuracy in genomic selection (Habier et al., 2007; Daetwyler et al., 2010). This pattern is also evident in ryegrass, as shown in Figure 6 under the parental model. Across all training–validation configurations, prediction accuracies were moderate to high and only approximately 10% lower than those obtained with a GBLUP model that exploits genomic information at the family rather than the parental level. In ryegrass, biparental crosses between single plants produce multiple F_1_ seeds that are subsequently multiplied in isolation, allowing intermating and resulting in the F_2_ seed used for field evaluation. Under this breeding scheme, Mendelian sampling effects are expected to largely cancel out when family means are considered (Daetwyler et al., 2013). The remaining differences in prediction accuracy may therefore be attributable to the larger number of genome-wide variants in the progeny-based model or to deviations from expected allele frequencies. Similar patterns have been reported in hybrid prediction studies in crops such as maize, rice, and sorghum, where genomic models based on parental information are able to predict cross performance with relatively high accuracy (Technow et al., 2014; Labroo et al., 2021; Sapkota et al., 2023). The genomic data used for prediction in this study also provided an opportunity to investigate the genetic architecture underlying forage quality traits.

The genetic diversity among selected parents and the high heritability for most of the traits allowed sufficient power to detect six QTL regions. Our results highlighted the importance of several SNPs on chromosome 1, with the most significant SNP for the NDFD trait located near predicted genes containing the homeobox (HD) domain, whose family encodes transcription factors involved in key biological processes such as meristem maintenance, embryonic patterning, lateral organ formation, and response to abiotic stress (Li et al., 2021). Studies in species such as rice, eucalyptus (Zhang et al., 2023), tomato (Li et al., 2021), banana, and apple demonstrate the central role of these genes in stress responses, reinforcing their relevance in plant adaptation to adverse environmental conditions. A study on gene expression in alfalfa (*Medicago sativa*) under thermal stress (Xu et al., 2023) observed the activation of HD genes in response to stress, suggesting their fundamental role in grasses as well, contributing to growth regulation and adaptation to environmental variations.

Two further markers–trait associations were detected. One is located near predicted genes with sequence homology to a 5’-3’ exonuclease domain, and another gene involved in glutamine metabolism. The exonuclease domain is linked to essential functions in DNA metabolism, including replication, repair, and recombination (Caponigro & Parker, 1995; Brune et al., 2005). The glutamine-related gene plays a crucial role in nitrogen metabolism, as glutamine is essential for the assimilation, transport, and redistribution of nitrogen in plants, while also being involved in the synthesis and regulation of amino acids, which are vital for proper growth and the nutritional content of plants (Yu et al., 2021; Nawaz et al., 2023).

Additional associations were detected at smaller peaks, including SNPs 3_159655121_C_T, 3_66395904_A_G, and 6_233416561_G_A. These regions are located near genes annotated as rhodanese-like and senescence-associated proteins, members of the NPF/peptide transporter family, and β-1,3-glucanases. These gene families have been implicated in nutrient remobilization during plant senescence, nitrogen transport and allocation, and carbohydrate metabolism and cell wall remodeling, respectively (Lim et al., 2007; Balasubramanian et al., 2012; Léran et al., 2014).

The QTLs identified in this study by back-solving GCA effects are promising for marker-assisted selection, whereas the flanking genes may be targets to better understand the expression of nutritive quality traits in perennial ryegrass, including digestibility. Improving such traits is not only essential for better animal performance, but also contributes to reducing methane emissions per unit of production, as forage quality and diet composition are key determinants of enteric methane intensity (Hristov, 2013; Galyon et al., 2026)

## Conclusions

Our results show that genomic-assisted prediction of single-cross performance is both feasible and accurate in perennial ryegrass when using a GCA model, despite its obligate outcrossing nature and family-level genetic heterogeneity. The limited genotype-by-environment and genotype-by-nitrogen-level interactions indicate that family performance was largely stable across these sources of variation and that genomic predictions are likely to be robust to nitrogen management and environmental conditions in Denmark. Furthermore, the identification of six QTL regions and plausible candidate genes highlights the value of leveraging existing genetic diversity in sparse crossing designs to dissect the genetic architecture of complex traits. From a practical breeding perspective, the parental model approach represents a cost-effective strategy, as it relies on genotyping parental lines rather than large numbers of progeny while still achieving competitive predictive accuracy.

## Data availability

All genotypic data (provided as a genomic relationship matrix) and phenotypic data used in this study are publicly available in the Dryad repository (https://doi.org/10.5061/dryad.q83bk3jm7).

## Acknowledgments

The authors acknowledge the financial support of the Danish Ministry of Food, Agriculture and Fisheries through the GUDP grant no. 34009-15-0952, which enabled the generation of the data used in this study.

## Conflicts of interest

The authors declare no competing interests.

## Notes

### Competing Interest Statement

The authors have declared no competing interest.

