## Supplementary Note S1 for "Genomic prediction of single cross families of perennial ryegrass in two nitrogen managements"

Prediction accuracies from the fitted parental models were on the high-end for most of the traits. A natural question that emerges is how such values compare to those obtained from a GBLUP model that takes a covariance matrix built with genome-wide variants from the $F_2$ families themselves. To answer that, we fitted the LMM model defined below in matrix notation:

$$\mathbf{y} = \mathbf{1}\mu + \mathbf{X_1 b_1} + \mathbf{X_2 b_2} + \mathbf{Z_1 u} + \mathbf{Z_2 u \cdot e} + \mathbf{Z_3 u \cdot n} + \epsilon$$

where, $\mathbf{y}$ is the vector of phenotypes; $\mu$ represents the overall intercept with associated vector $\mathbf{1}$ containing 1's; $\mathbf{b_1}$ is a vector of fixed effects of environment and nitrogen management classes, $\mathbf{b_2}$ is the fixed effect for classes created by combine harvesters used and the date of harvest; $\mathbf{X_1}$ and $\mathbf{X_2}$ are the associated incidence matrices for the fixed effects, respectively; $\mathbf{u}$ is the family random effect assumed $\mathbf{u} \sim N(\mathbf{0}, \mathbf{G} \cdot \sigma^2_u)$; $\mathbf{u \cdot e}$ is a vector of random family by environment interaction effects, following $\mathbf{u \cdot e} \sim N(\mathbf{0}, \mathbf{G} \cdot \sigma^2_{u \cdot e})$; $\mathbf{u \cdot n}$ is a vector of the random interaction effect of family by nitrogen, assumed $\mathbf{u \cdot n} \sim N(\mathbf{0}, \mathbf{G} \cdot \sigma^2_{u \cdot n})$; $\epsilon$ is the vector of random residual effect, which follows $\epsilon \sim N(\mathbf{0}, \mathbf{I} \cdot \sigma^2_\epsilon)$; $\mathbf{Z_1}$, $\mathbf{Z_2}$, and $\mathbf{Z_3}$ are incidence matrices linking phenotypic observations to the respective random effects, and $\mathbf{I}$ refers to an identity matrix. The genomic relationship matrix $\mathbf{G}$ for the F2 families was computed using allele frequencies and a total of 56,645 filtered variants (see Bornhofen et al., 2023 for genotyping details and $\mathbf{G}$ construction). Model comparison was performed using 5-fold cross-validation (CV), repeated 10 times, as described in the Materials and Methods section, and the results are shown in Figure S1.

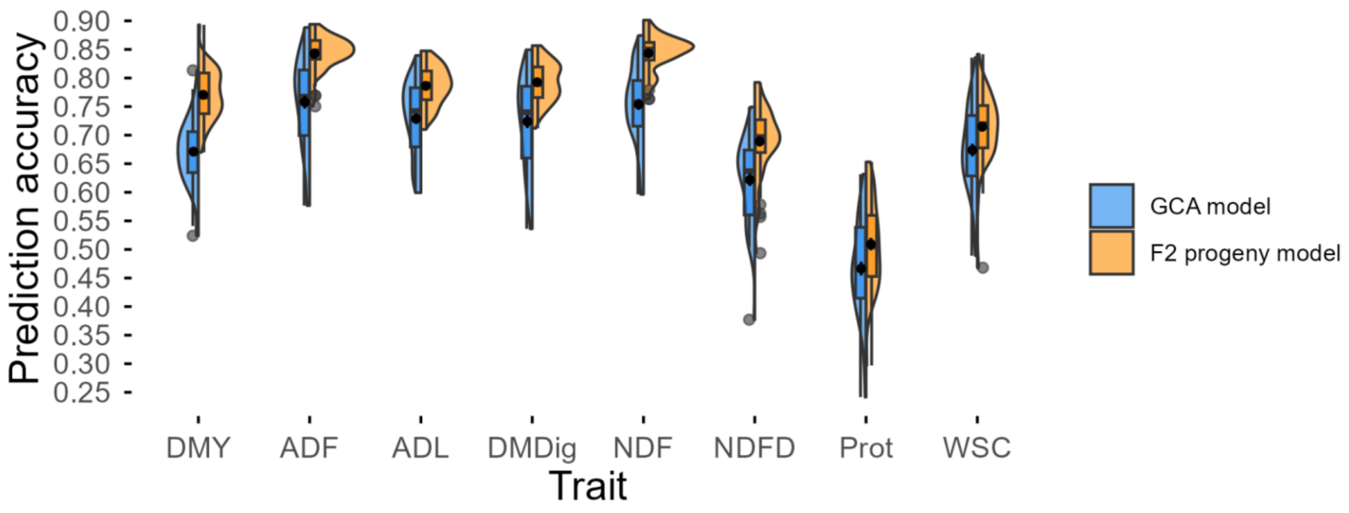


**Figure S1.** Genomic prediction using variant information at the progeny level yields on average around 10% gain in prediction accuracy over the general combining ability (GCA) model.

Accuracies were computed as the Pearson correlation coefficient between predicted family breeding values from a repeated 5-fold cross-validation scheme and adjusted means.
